# Elevational and latitudinal range extensions of the puna mouse, *Punomys* (Cricetidae), with the first record of the genus from Chile

**DOI:** 10.1101/2023.03.07.531530

**Authors:** Marcial Quiroga-Carmona, Jay F. Storz, Guillermo D’Elía

## Abstract

We report an elevational record for the Andean sigmodontine puna mouse *Punomys*, which is also the first record of the genus in Chile. The record is based on a mummified specimen that we discovered at an elevation of 5,461 m (17,917’) in the caldera of Volcán Acamarachi, Región de Antofagasta, Chile. Results of a morphological suggest that the specimen can be provisionally referred to the species *P. lemminus*. This new record also extends the known geographic distribution of the genus by 700 km to the south and brings the known Chilean mammal richness to a total of 170 living species and 88 genera. This finding highlights the need for increased survey efforts in more remote, high-elevation regions and demonstrates that there is still much to be learned about the mammal fauna of the Andean Altiplano.

## INTRODUCTION

The Linnaean and Wallacean shortfalls refer to the discrepancy between known and actual numbers of species and the discrepancy between the known and actual range limits of species, respectively (Hortal et al. 2015). Reducing these knowledge gaps requires specimen collection in the field, especially in remote regions of the planet that are still comparatively underexplored by biologists (Patterson 2002). In spite of more than two centuries of zoological research in South America, mammal assemblages from large portions of the continent remain poorly characterized, and this is especially true for small mammal assemblages in the Andean Altiplano.

Puna mice of the genus *Punomys*, Osgood, 1943, represent a particularly enigmatic group of Andean sigmodontine rodents. The genus comprises two recognized species, *P. lemminus* Osgood, 1943, and *P. kofordi* Pacheco and Patton, 1995. These medium-sized, stoutly built mice have a vole-like appearance with long and lax fur, a short tail, and relatively reduced ears (Patton 2015). All known *Punomys* specimens have been collected from a small number of localities in the Altiplano of southern Peru and northwestern Bolivia, at elevations between 4,550 and 4,770 m (see Map 44 from Patton et al. 2015). The two described species of *Punomys* differ in body proportions, pelage color (including dorso-ventral contrast), and several craniodental traits, such as convergence of the zygomatic arches, narrowing of the zygomatic plate, breadth of the nasal region, and size of procingular conules of upper molars (see Pacheco and Patton 1995; Patton 2015). However, the so far last study centered on *Punomys*, suggested that the distinction between both of its nominal forms need to be better evaluated (Salazar-Bravo et al. 2011).

In recent years, high-elevation surveys in the Central Andes have revealed how much we have yet to learn about the elevational range limits of small mammals. For example, mountaineering mammal surveys of multiple volcanoes in the Chilean Puna de Atacama have documented evidence of the Andean leaf-eared mice, *Phyllotis vaccarum* (previously referred as to *P. xanthopygus rupestris*), living at elevations >6000 m (Storz et al. 2020; Steppan et al. 2022), and mammal surveys in the Andean Altiplano of northwestern Argentina have documented surprisingly diverse rodent assemblages at elevations >4000 m (Urquizo et al. 2022). Thus, as part of a continuing effort to further redress Linnean and Wallacean shortfalls for Andean small mammals, here we report the first Chilean record of the genus *Punomys*, a record that also significantly extends the elevational and latitudinal range of *Punomys*.

## MATERIALS AND METHODS

### Specimen collection, preparation, and morphological characterization

During the course of a high-elevation mammal survey in the dry Puna ecoregion of the Chilean Andes, we discovered a mummified rodent carcass in the caldera of Volcán Acamarachi, Region de Antofagasta (23°17’33.88” S, 67°37’4.48” W; Figure 1). We deposited the specimen (Field number: MQC 385) in the Colección de Mamíferos, Universidad Austral de Chile (UACH) (catalog number UACH 8478). The mummy was completely intact, with the tail, and fore- and hindfeet firmly attached to the body, and pelage in good condition. After taking several photographs to record the external appearance of the specimen, we removed the skull and mandible to assess craniodental traits to compare them with the descriptions of the species of *Punomys* (e.g., Osgood 1943; Pacheco and Patton 1995; Patton 2015). With the same purpose, we also recorded 13 cranial measurements with a digital caliper to the nearest 0.01 mm, following the protocol of Salazar-Bravo et al. (2011).

**Figure 1.**
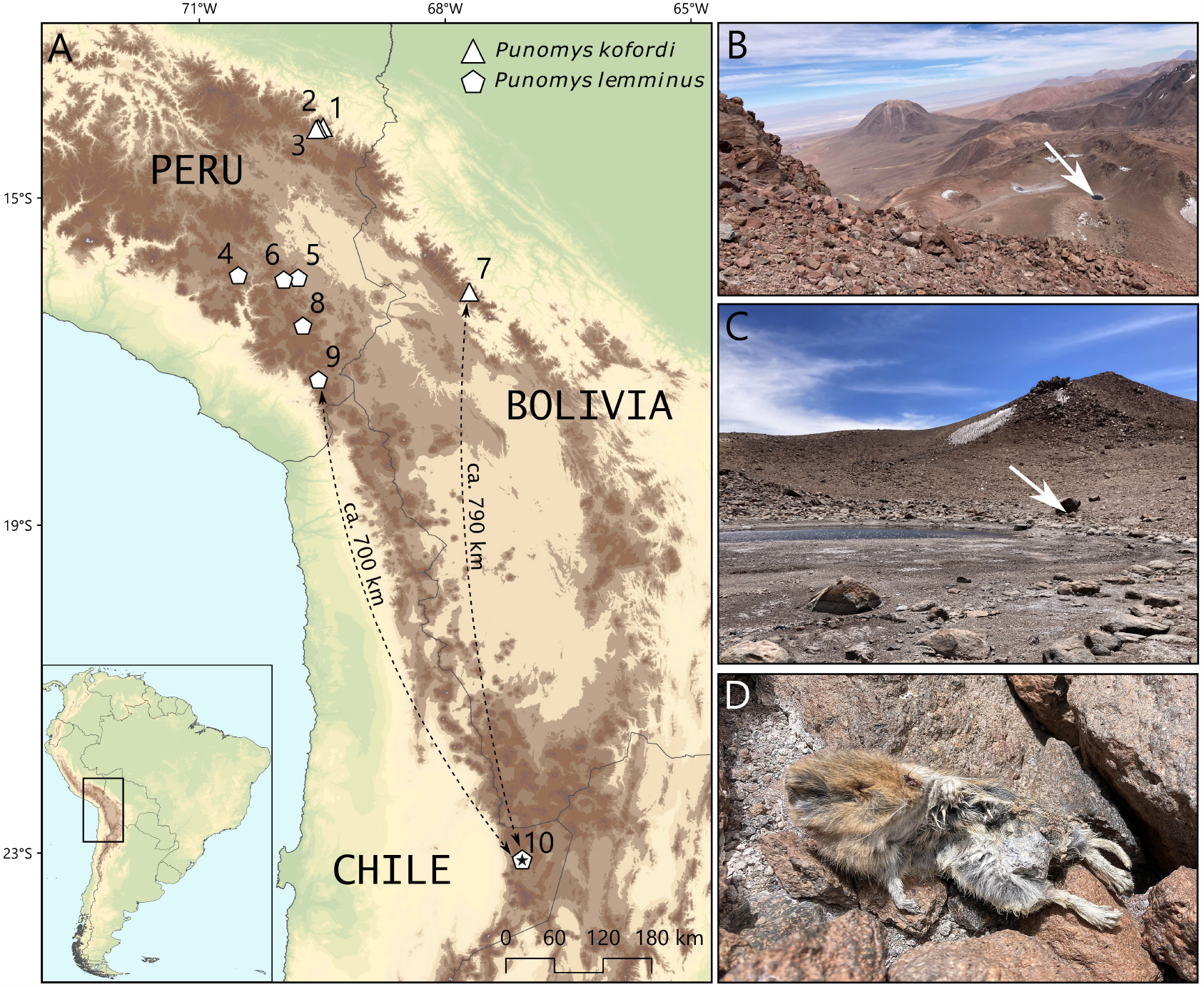
(A) Recording localities of *Punomys* including the locality where the mummy specimen was discovered in the caldera of Volcán Acamarachi, Chile (pentagon with black star). Numbers correspond to localities listed in Supplementary material 1. Brown-shading denotes Altiplano terrain >4,000 m. (B) Aerial view of the locality where the mummy of *Punomys* was discovered at an elevation of 5,461 m in the caldera of Volcán Acamarachi, Región de Antofagasta, Chile (23°16.98’ S, 67°37.46’ W). The view is from the summit of Acamarachi (6,046 m [19836’]) on the southern rim of the caldera. The mummy of *Punomys* was found at the base of a boulder near the edge of a small glacial pond (denoted by arrow). (C) Close-up view of the edge of the glacial pond and the boulder (denoted by arrow) where the mummy was found. (D) Mummy of *Punomys* at the site of discovery.

### Tissue sampling and DNA sequencing

After removing the skull, we preserved muscle tissue samples in ethanol as a source of genomic DNA. After extracting DNA, we PCR-amplified a 801 bp fragment of the mitochondrial gene, *cytochrome-b* (*Cytb*), using primers MVZ 05 and MVZ 16 (da Silva and Patton 1993) in accordance with the protocol of Teta et al. (2013).

Amplicons were purified and sequenced by Macrogen Inc., Korea. Since in GenBank the genus *Punomys* is only represented by two *Cytb* sequences (JQ434426 of 1139 bp and KY754123 of 426 bp) which were generated from a single specimen (VPT1890) of *P. kofordi*, we obtained additional tissue loans from a total of six *Punomys* specimens housed in the Museum of Vertebrate Zoology (MVZ), University of California-Berkeley (*P. kofordi*: MVZ114757, MVZ114758, MVZ116190, MVZ139589; *P. lemminus*: MVZ115948, MVZ116036). For each of these six specimens, we extracted genomic DNA from toe pads of museum skins. We performed the PCR-amplification of *Cytb* using the internal primer Oct349, following the protocol of Cadenillas and D’Elía (2021), and amplicons were purified and sequenced as described above. However, given the degradation levels of these samples, sequences of sufficient quality were recovered for only four specimens (MVZ114757, MVZ115948, MVZ116036, and MVZ139589), which were included in subsequent analyses. We deposited all new sequences in GenBank under accession numbers OQ571714-OQ571718 (see Supplementary material 1).

We created a *Cytb* sequence matrix that included all newly generated and previously generated *Punomys* sequences as well as sequences from representatives of the four main phylogroups of *Andinomys edax* (Jayat et al. 2017; KY608037, KY608044, KY608047, KY608056). The monotypic genus *Andinomys* is the sister group of the genus *Punomys* (Salazar-Bravo et al. 2013), and the *Andinomys*-*Punomys* clade is recognized as the tribe Andinomyini. A recent phylogenomic analysis Parada et al. (2021) found that Andinomyini is sister to a clade form by the tribes Abrotrichini, Euneomyini and Phyllotini. We therefore used representative *Cytb* haplotypes from each of these tribes (MN275227 *Abrothrix longipilis*; AY95623 *Phyllotis darwini*; HM167872 *Euneomys chinchilloides*) to root the tree.

### Phylogenetic analysis

We aligned *Cytb* sequences using MAFFT v7 (Katoh et al. 2017) under E-INS-i strategy to establish character primary homology. We then visually inspected the alignment with AliView v1.26 (Larsson 2014) to check for the presence of internal stop codons and shifts in the reading frame. JModelTest2 (Darriba et al. 2012) was employed to identify the TIM2+F+G4 as the best nucleotide substitution model for the dataset, based on the Bayesian Information Criterion (BIC). We estimated a *Cytb* gene tree using a Bayesian Inference (BI) phylogenetic analysis implemented in BEAST 2, version 2.6.2 (Bouckaert et al. 2014), running 100 × 10^6^ generations, and sampling produced trees every 1000 generations. However, as this model cannot be implemented in BEAST2, we employed the GTR+G+I model, according to Ronquist and Huelsenbeck (2003). The first 10% of the total trees were discarded and the remaining trees were used to construct a maximum clade credibility tree with posteriori probability values (PP), employing TreeAnnotator version 2.6.0 (Rambaut and Drummond 2019). Finally, we estimated the percent of sequence divergence between haplotype pairs of *Punomys* using MEGA X 10.1.8 (Kumar et al. 2018) in the form of p-distances, ignoring those sites with missing data.

## RESULTS

The Chilean specimen of *Punomys* (UACH 8478) was found at an elevation of 5,461 m (17,917’) within the caldera of Volcán Acamarachi (also known as Cerro Pili). This places is devoid of vegetation and has a substrate of volcanic rock and sand, which continues with a steep rocky couloir that leads to the summit at 6046 m asl (19,836’; see Figure 1B-C). The suite of characteristics of the external morphology of the mummified specimen that could be examined, conforms to the diagnostic criteria described by Patton (2015) for *Punomys*. Hindfoot soles are proximally hairy and with well-defined pads; toes are of medium length with short claws. Not all external morphological characters (e.g., length of ears, legs and body) could be properly measured on the mummy specimen due to its preservation condition. However, in spite of slight discoloration, the dorso-ventral contrast of the general pelage is clear in this specimen and conforms to that of *P. lemminus*. This contrast is most conspicuous in the pelage of the cheeks and throat (Figure 2).

**Figure 2.**
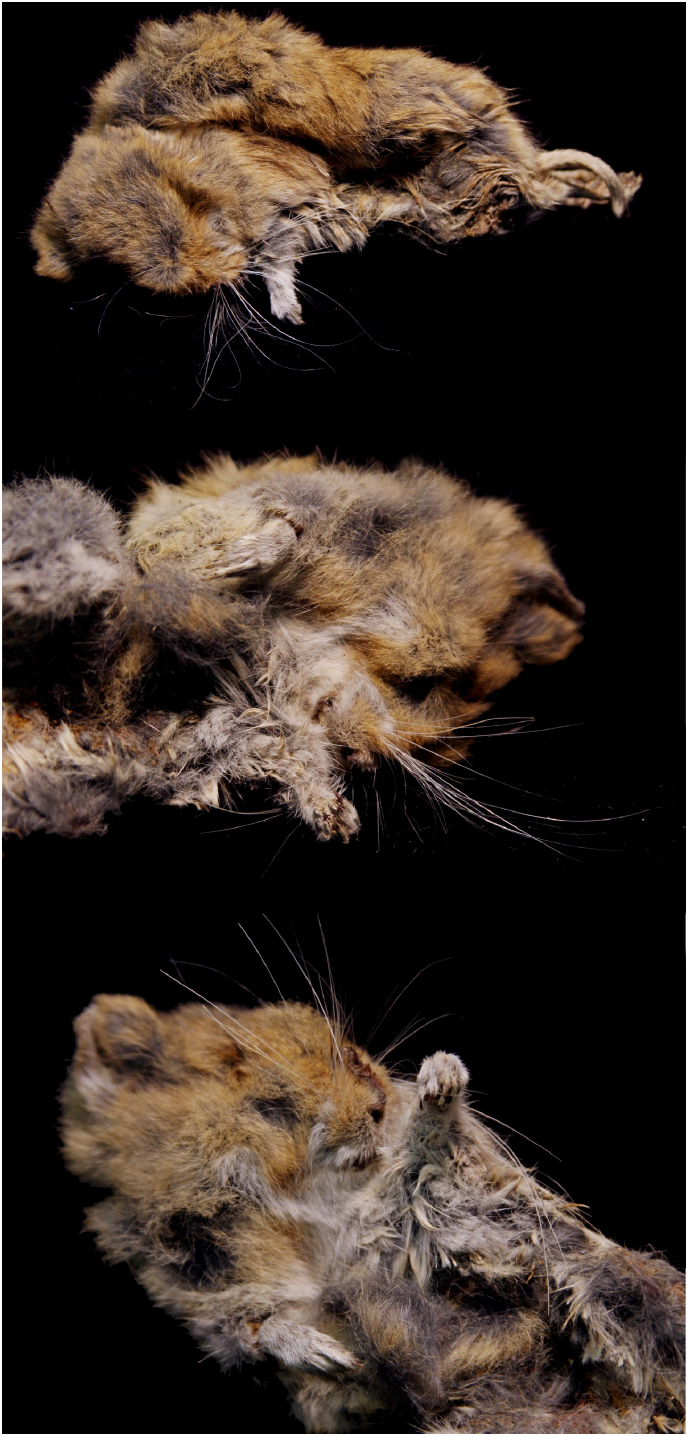
External views of the mummified specimen of *Punomys* from Volcán Acamarachi, Chile (UACH 8478). From top to bottom, dorsal general appearance of pelage, and close-up of the right and left sides of the face, throat and chest of the mummy, showing the dorso-ventral contrast of fur coloration.

The skull of the Chilean specimen is robust and heavy, with a broad and short rostrum and a massive and squared braincase (Figure 3). The most noteworthy cranial features include broad and anteriorly expanded nasals, narrow interorbital region, robust zygomatic arches, mesopterygoid fossa extending near to posterior edge of M3, and large and inflated tympanic bullae, which together are part of the diagnostic state characters of *Punomys* (Patton 2015).

**Figure 3.**
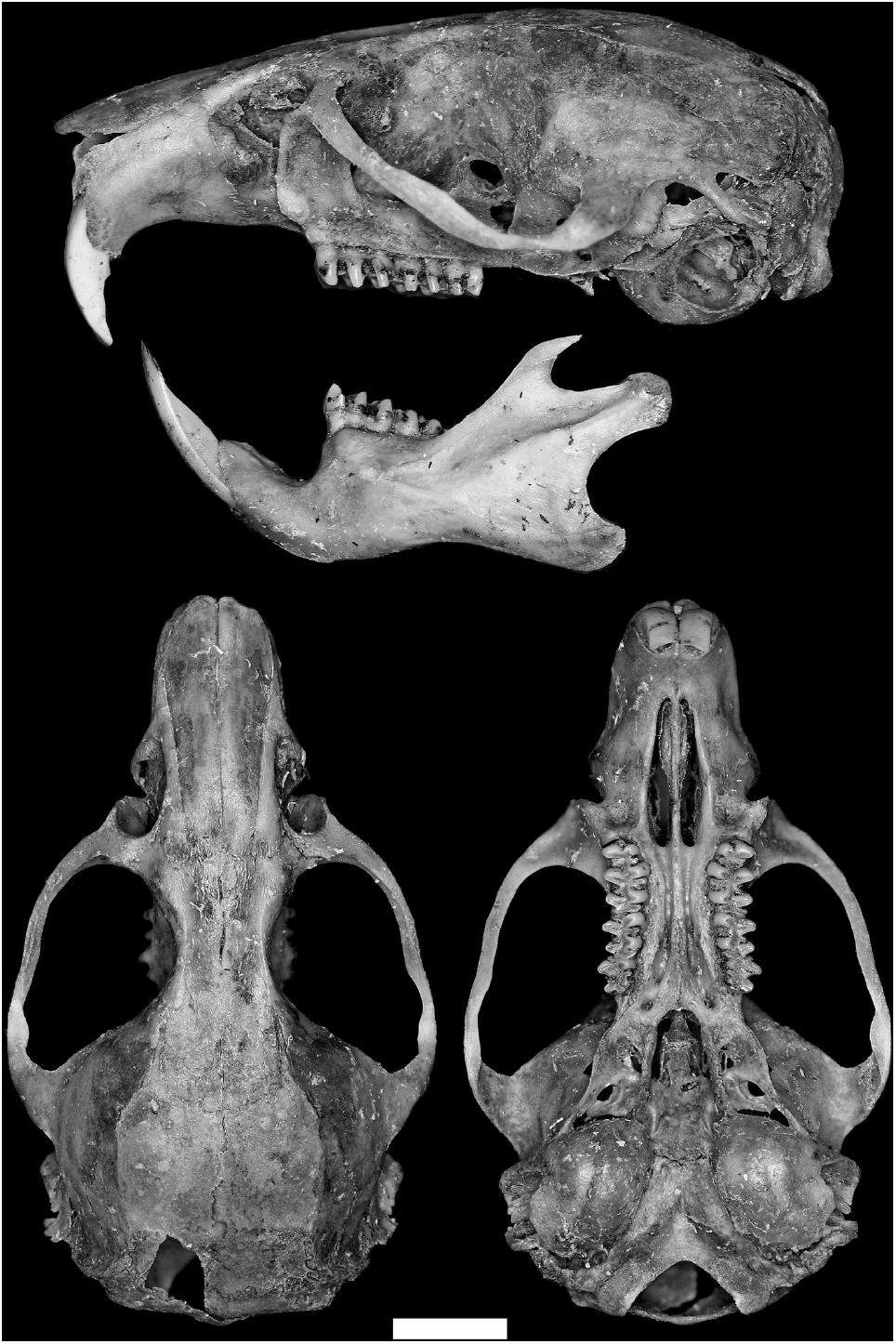
Skull and jaw of the specimen of *Punomys* from Volcán Acamarachi, Chile (UACH 8478). Scale bar = 5 mm.

Measurements of each cranial character of the Chilean specimen (Table 1) fall within the range of values reported for the genus *Punomys* (see Pacheco and Patton 1995; Salazar-Bravo et al. 2011), but do not permit conclusive inference about species identity as some measurements were more similar to those reported for *P. lemminus* (e.g., incisive foramen length, palatal bridge length) and others were more similar to those reported for *P. kofordi* (e.g., greatest length of skull, condyloincisive length). Comparative examination of skull photographs of the holotype of *P. kofordi* (MVZ 139588) and reference material of *P. lemminus* (MVZ 115948) presented by Pacheco and Patton (1995) reveals that the convergent shape of the zygomatic arches and the presence of a deep zygomatic notch of the Chilean specimen conforms to the morphotype of *P. lemminus* (see Figs. 2 and 3 in Pacheco and Patton 1995). Dental characters of the Chilean specimen are also characteristic of *Punomys* (Pacheco and Patton 1995, Patton 2015): molars are complex, with the presence of a lingual style more developed that the labial style, M1 are curved and with the procingulum anteriorly divergent, with anterolabial and anterolingual conules with subequal sizes, and M3 have four well-defined cusps (Figure 4).

**Table 1.** Cranial measurements of specimens of *Punomys kofordi* and *P. lemminus*, the Bolivian specimen of *P. kofordi* reported by Salazar-Bravo et al. (2011, CBF 858), and the mummified specimen from Volcán Acamarachi, Chile, UACH 8478. Mean and range of the presented measures correspond to those described by Pacheco and Patton (1995) and by Salazar-Bravo et al. (2011), for Peruvian and Bolivian specimens. All measurements are in millimeters.

| *P. kofordi*<br>(n = 6)<br>Pacheco and Patton<br>(1995) | *P. kofordi*<br>(n = 3)<br>Salazar-Bravo<br>et al. (2011) | *P. kofordi*<br>CBF 858<br>Salazar-Bravo<br>et al. (2011) | *P. lemminus*<br>(n = 3)<br>Pacheco and Patton<br>(1995) | *Punomys* Chile<br>UACH 8478<br>This study |
| --- | --- | --- | --- | --- |
| Greatest length of skull<br>32.9 (31.4-33.9) | 35.2 (35.1-35.4) | -- | 33.3 (32.5-34.1) | 34.32 |
| Condylloincisive length<br>30.9 (28.8-31.9) | 33.7 (33.3-34.3) | -- | 31.8 (31.0-32.2) | 33.40 |
| Zygomatic breadth<br>17.6 (16.2-18.8) | 18.4 (18.4-18.4) | -- | 18.9 (18.6-19.3) | 20.22 |
| Braincase breadth<br>14.7 (14.5-15.2) | 13.8 (13.1-14.6) | 13.6 | 15.2 (14.7-15.5) | 15.27 |
| Interorbital breadth<br>4.4 (4.3-4.5) | 4.5 (4.4-4.7) | -- | 4.2 (4.0-4.3) | 4.53 |
| Diastema length<br>8.0 (7.4-8.4) | 9.0 (8.7-9.3) | 8.5 | 8.3 (8.0-8.7) | 9.08 |
| Maxillary tooth row length<br>6.9 (6.7-7.2) | 6.7 (6.6-6.9) | 6.7 | 7.0 (6.5-7.5) | 7.36 |
| Incisive foramen length<br>7.0 (6.9-7.2) | 8.1 (7.8-8.5) | 7.5 | 6.8 (6.7-6.9) | 6.96 |
| Palatal bridge length<br>6.6 (6.4-6.8) | -- | -- | 7.4 (6.5-8.0) | 7.76 |
| Rostral breadth<br>6.9 (6.4-7.3) | 6.6 (6.5-6.7) | 6.6 | 7.2 (7.0-7.6) | 6.41 |
| Palatal bridge width<br>2.3 (1.8-2.9) | 2.5 (2.48-2.58) | 1.9 | 2.2 (2.1-2.3) | 2.61 |
| First upper molar breadth<br>2.1 (2.0-2.2) | -- | -- | 2.1 (2.0-2.2) | 2.16 |
| Zygomatic plate breadth<br>3.2 (2.8-3.4) | 3.6 (3.5-3.7) | 2.9 | 3.6 (3.5-3.6) | 3.53 |

**Figure 4.**
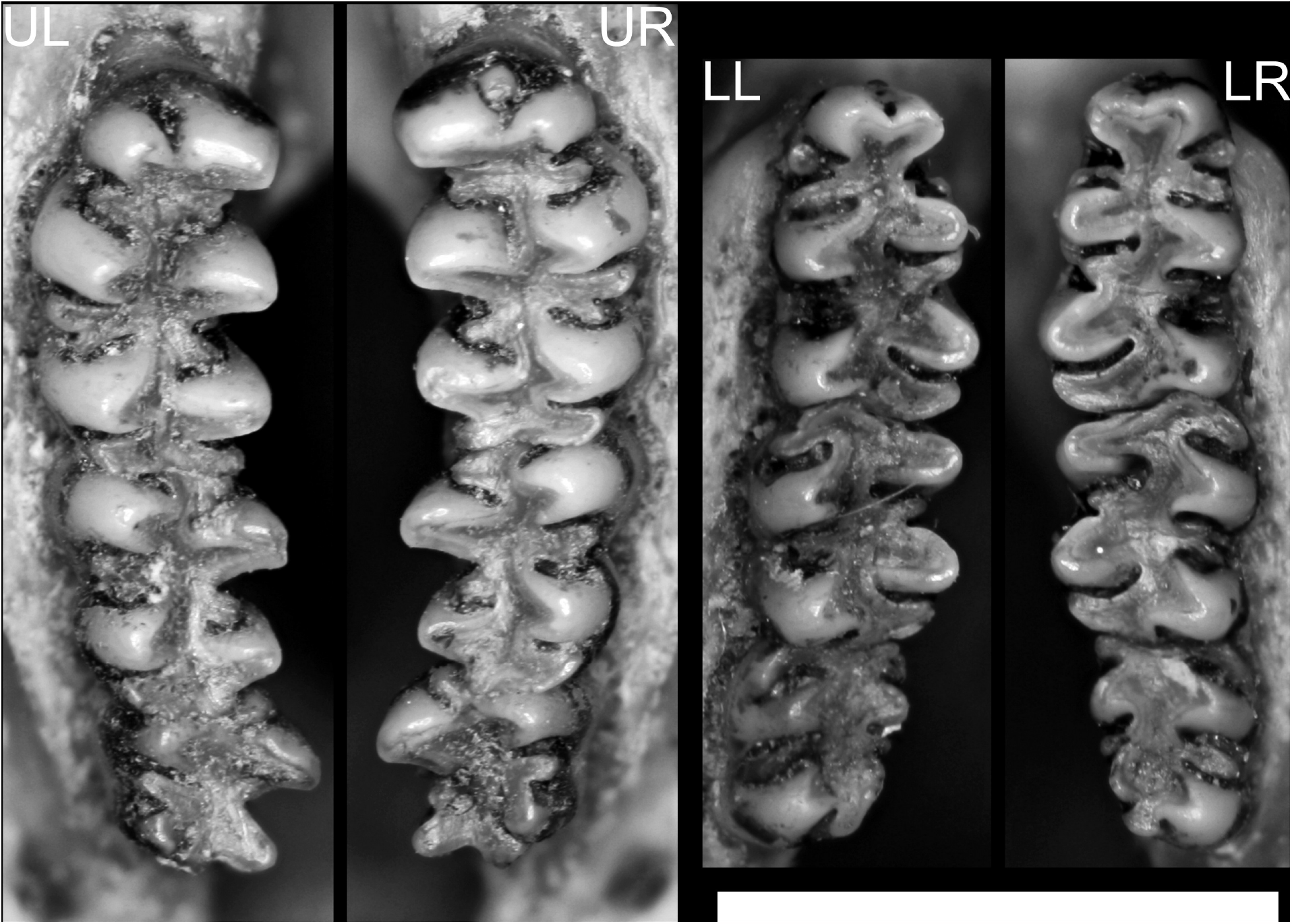
Occlusal views of the molar series of the specimen of *Punomys* from Volcán Acamarachi, Chile (UACH 8478). U and L denote upper and lower rows, and L and R to left and right ones, respectively. Scale bar = 5 mm.

Phylogenetic analysis of *Cytb* sequences (Figure 5) revealed that the Chilean specimen groups with the other sequences of *Punomys* in a well-supported clade (Bayesian posterior probability: PP = 1). Within this clade, haplotypes representing the nominal species *P. kofordi* and *P. lemminus* did not form reciprocally monophyletic groups and the Chilean specimen (UACH 8478) is recovered as sister to a haplotype of a specimen MVZ 139589 assigned to *P. kofordi* and collected at a locality close to the type locality of this form (Figure 1; Supplementary Material 1). Pairwise genetic distances between the Chilean specimen and all other *Punomys* specimens ranged from 1.55% to 3.12%, while pairwise comparisons for the whole sample of *Punomys* ranges between 0.23% and 3.12% (Table 2).

**Table 2.**
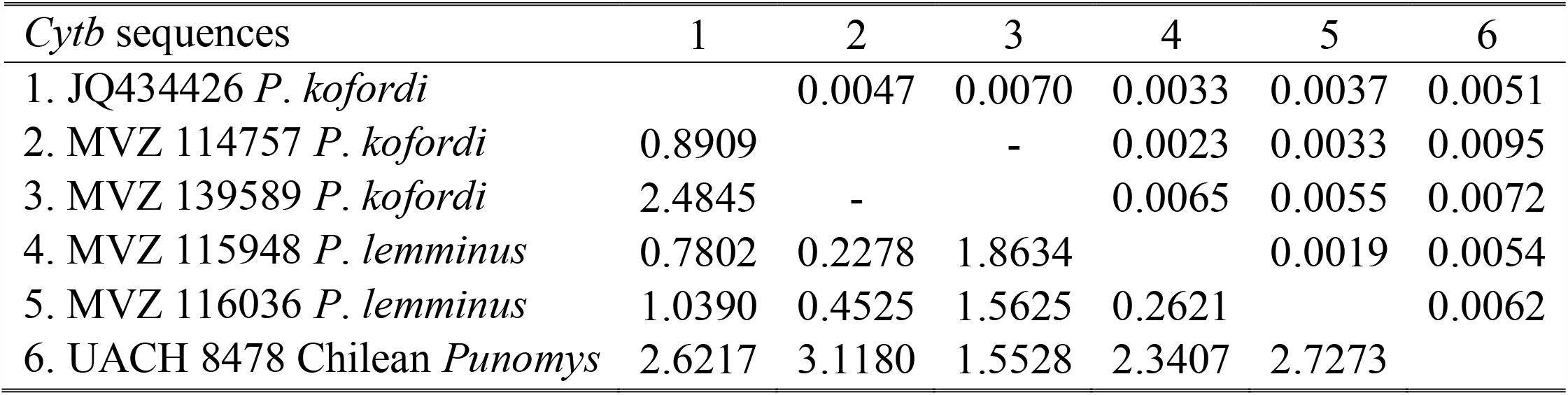
Percentage of the observed genetic divergence between *Cytb* haplotype pairs retrieved from specimens of *Punomys*. Standard error estimate(s) are shown above the diagonal. Comparison between sequences MVZ 114757 and MVZ 139589 could not be done due to the lack of bases in common between both sequences

**Figure 5.**
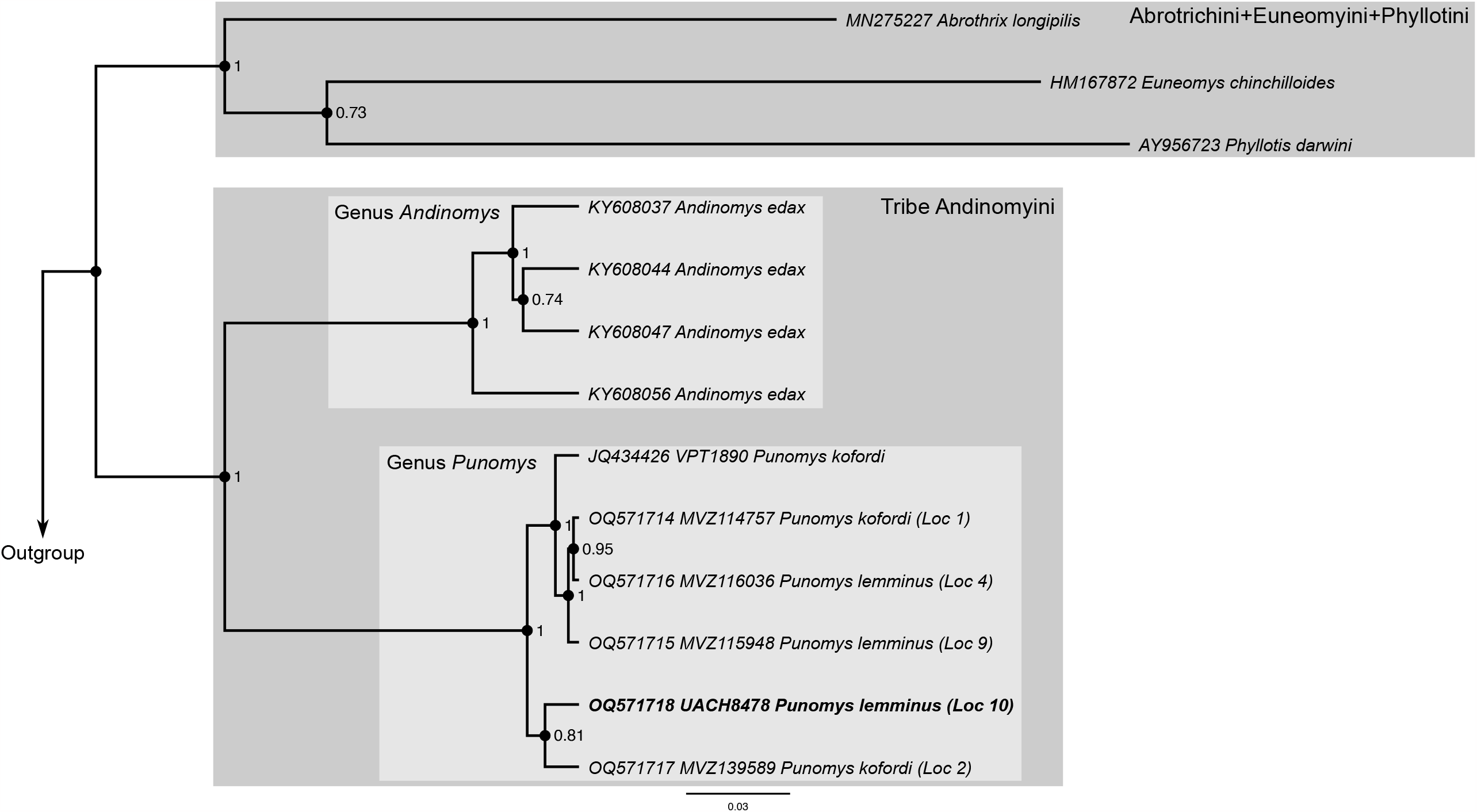
Bayesian inference tree based on *Cyt* B gene variation showing the placement of the haplotype recovered from the *Punomys* specimen from Volcán Acamarachi, Chile (UACH 8478). Numbers denote posterior probability values for the adjacent nodes; only values for species clades and relationships among them are shown. For representative haplotypes of *Punomys*, GenBank accession number, museum catalog number, and locality according to Figure 1 are provided.

## DISCUSSION

Analysis of morphological characters and *Cytb* sequences clearly demonstrate that the mummy specimen from Volcán Acamarachi (UACH 8478) is referable to *Punomys*, making it the first record for this genus from Chile. Given that we discovered the mummy at an elevation of 5,461 m, the Chilean specimen also represents an elevational record for the genus *Punomys*, as previous specimens in Peru and Bolivia were collected between 4,450 and 4,770 m asl (Pacheco and Patton 1995; Salazar-Bravo et al. 2011). With this new specimen-based elevational record, *Punomys* is second only to *Phyllotis vaccarum* as the mammal taxon with the highest elevational range limit (Storz et al. 2020). In addition to the new elevational record for *Punomys*, our Chilean specimen also represents a ∼700 km southward extension of the known latitudinal range of the genus. Our discovery of the Chilean specimen at such an extreme elevation in the Central Volcanic Zone suggests that the taxon could be continuously distributed along the upper reaches of the Andean Cordillera in northern Chile and bordering regions with Peru and Bolivia, and likely also Argentina (Figure 1). This is a testable hypothesis that requires more survey efforts at extreme elevations.

Although specimen UACH 8478 is clearly referable to the genus *Punomys*, it is difficult to make a species-level determination, largely because diagnostic differences between *P. kofordi* and *P. lemminus* are not clearly defined (Salazar-Bravo et al. 2011). The lack of reciprocal monophyly between *Cytb* haplotypes of the two nominal species reinforces the notion that the species boundaries of *Punomys* must be reviewed. Thus, operating under the assumption that these nominal forms represent distinct species, three morphological characters appear to be especially informative with regard to the identity of the Chilean specimen: the dorso-ventral contrast of fur coloration, convergent shape of the zygomatic arches, and the deepest of the zygomatic notch (Pacheco and Patton 1995), provide a basis for provisionally recognizing the Chilean *Punomys* specimen as *P. lemminus*. Notwithstanding, metric characters such as greatest length of skull and condyloincisive length also suggest some to affinity with *P. kofordi*, but considering the small size of the available samples this can be ruled out. On this regard, even when the haplotype of the Chilean specimen appears as sister to a haplotype of *P. kofordi*, this relationship lacks significant support (PP = 0.81; Figure 5) and the results of phylogenetic analysis are not informative on to which species the Chilean specimen belongs since these two nominal forms are not monophyletic.

The provisional designation of Chilean *Punomys* as *P. lemminus* could makes biogeographic sense, given that Volcán Acamarachi and the Peruvian collection localities for *P. lemminus* are situated in the western slopes of the Andes, and are therefore connected by a continuous tract of highland terrain, whereas the collection localities for *P. kofordi* lie much further to the north in the eastern side of the Andes (Figure 1A). However, biogeographic plausibility by itself does not provide a reliable means of taxonomic identification. In summary, the clarifying of the taxonomic identity of the Chilean specimen requires an assessment of the species limits within *Punomys*, which in turn needs the collection of additional specimens.

Our discovery highlights the value of conducting surveys in inaccessible, poorly explored regions of South America, and suggests that the Andean Altiplano is a region where much work is still needed to reduce the Linnean and Wallacean shortfalls for native small mammals (Storz et al. 2020; Rengifo et al. 2022). With our new record of *Punomys*, the number of living mammal species and genera in Chile now stands at 170 and 88, respectively (D’Elía et al. 2020; Rodríguez-San Pedro et al. 2022, 2023; Quiroga-Carmona et al. 2022). Recent additions to the Chilean mastofauna have been based on descriptions of new species (e.g., *Abrothrix manni* and *Dromiciops bozinovici* and *D. mondaca*; D’Elía et al. 2015, 2016a), as well as range extensions for previously described species (e.g., *Eumops perotis, Histiotus laephotis, Notiomys edwardsii, Nyctinomops aurispinosus, Oligoryzomys flavescens*; Ossa et al. 2015; D’Elía et al. 2016b; Rodríguez-San Pedro et al. 2022, 2023; Quiroga-Carmona et al. 2022). More field-based specimen collection is needed to further redress Linnean and Wallacean shortfalls in our knowledge of biodiversity in South America and elsewhere. The required efforts would be facilitated by simplifying permitting procedures to conduct biological surveys (D’Elía et al. 2019, Alexander et al. 2021, Teta 2021).

## ACKNOWLEDGEMENTS

We thank Jim Patton and Chris Conroy (Museum of Vertebrate Zoology) for specimen loans, Alex González and Chandrasekhar Natarajan for assistance with laboratory work, and Mario Pérez Mamani and Juan Carlos Briceño for assistance and companionship in the field.

## Author contributions

MQC and JFS performed field work. MQC and GD generated and analyzed data and wrote the manuscript. All authors have read, commented, and agreed to the final draft of the manuscript.

## Research funding

NSF Grants IOS-2114465, OIA-2140377, and OIA-2140307, National Geographic Explorer Grant NGS-68495R-20, FONDECYT Grant 1221115.

## Conflict of interest statement

The authors declare no conflicts of interest regarding this article.

## SUPPLEMENTARY MATERIAL 1. Localities where specimens of *Punomys* have been collected and material housed in biological collections

Localities are plotted in Figure 1, corresponding to the records of *Punomys* along the Central Andes, in Bolivia, Chile and Peru. Localities are ordered by species and latitudinally from north to south. Collection numbers of the specimens collected at each locality are provided. In addition, Genbank accession numbers of the five *Cytb* sequences generated during this study are here provided (OQ571714-OQ571718).

*Punomys kofordi* Osgood, 1943

Locality 1. PERU, Puno, 8 mi SSW Limbani; coordinates: 14°15.30′S, 69°44.04′W. Material: MVZ 114757 (OQ571714) -114759, 116190-116195 (Paratypes). See locality 7 in the Figure 1 of Pacheco and Patton (1995).

Locality 2. PERU, Puno, 15 mi ENE Crucero, Aricoma Pass; Coordinates: 14°16.02′S, 69°47.64′W. Material: MVZ 139589 (OQ571717). See locality 6 in the Figure 1 of Pacheco and Patton (1995).

Locality 3. PERU, Puno, Lago Aricoma, 13 mi ENE Crucero, Abra Aricoma; coordinates: 14°16.68′S, 69°49.32′W. Material: MVZ 139588 (Holotype). See locality 5 in the Figure 1 of Pacheco and Patton (1995).

*Punomys lemminus* Pacheco and Patton, 1995

Locality 4. PERU, Arequipa, Huaylarco, 55 mi ENE Arequipa; coordinates: 16°01.98′S, 70°49.98′W. Material: MVZ 116036 (OQ571716). See locality 1 in the Figure 1 of Pacheco and Patton (1995).

Locality 5. PERU, Puno, west of Ilave; coordinates: 16°05.40′S, 70°06.12′W. Material: MCZ 42883. See MCZ database https://mczbase.mcz.harvard.edu/Specimens.cfm.

Locality 6. PERU, Puno, San Antonio de Esquilache; coordinates: 16°06.36’S, 70°16.80′W. Material: FMNH 49710 (holotype). See locality 2 in the Figure 1 of Pacheco and Patton (1995).

Locality 7. BOLIVIA, La Paz, Cumbre del camino a Yungas; coordinates: 16°19.80′S, 68°01.20′W. Material: CBF 858. See Figure 1 in Salazar-Bravo et al. (2011).

Locality 8. PERU, Puno, Caccachara, about 5 mi SW crest of the western Cordillera, approximately 50 mi SW Ilave; coordinates: 16°40.98′S, 70°04.02′W. Material: MCZ 42789-4291. See locality 3 in the Figure 1 of Pacheco and Patton (1995).

Locality 9. PERU, Tacna, 20 km NE Tarata; coordinates: 17°20.82′S, 69°54.00′W. Material: MVZ 115948 (OQ571715). See locality 4 in the Figure 1 of Pacheco and Patton (1995).

Locality 10. CHILE, Región de Antofagasta, Comuna de San Pedro de Atacama, southeast flank of Volcán Acamarachi. Elevation: 5461 m asl. Coordinates: 23°17.58′S, 67°37.08′W. Material: UACH 8478 (MQC 358; OQ571718).

## Notes

### Competing Interest Statement

The authors have declared no competing interest.

